# Evaluation of Field Sampling Techniques for Environmental Microbial Exposure: Assessing Efficacy and Feasibility

**DOI:** 10.1101/2020.06.14.150722

**Authors:** Kathryn R. Dalton, Kristoffer Spicer, Shanna Ludwig, Dorothy Clemons-Erby, Timothy Green, Ana M. Rule, Kirsten Koehler, Meredith C. McCormack, Meghan F. Davis

**Author notes:** Corresponding author: Kathryn R. Dalton, Johns Hopkins University Bloomberg School of Public Health, 615 N Wolfe St, EHE W7034G, Baltimore MD 21205. **Disclosure Statement:** None of the authors of this manuscript has a financial interest or benefit arising from the direct applications of this research. No specific funding was received for this project.

## Abstract

Environmental exposures in schools, including microbial exposures, can lead to detrimental childhood health outcomes. We evaluated two sampling techniques – standard flocked swabs versus sterile electrostatic cloths – to quantify *Staphylococcus* bacterial burden from school surfaces. Electrostatic cloths demonstrated higher test sensitivity and yielded higher surface area-standardized colony forming units compared to swabs. Despite protocol standardization, consistently larger surface areas were sampled with electrostatic cloths. This suggest that electrostatic cloths were more effective and practical for fieldwork.

## Introduction

Sampling protocols in environmental exposure assessment studies are a critical part of the overall research design, yet frequently receive minimal attention and critique. More commonly, sampling protocols are selected based on previously validated studies without consideration to their utilization in a novel field setting. This project originated at the appeal of field research technicians, who wanted to adopt a sampling protocol previously validated in households, for use in a classroom setting ^1^. This work was part of larger study evaluating physical, chemical and biological (allergen and microbial) environmental exposures in inner-city schools and their association to childhood health and academic performance ^2,3^.

Environmental microbial exposures—specifically exposures to the common opportunistic pathogen *Staphylococcus aureus*—have been associated with both infectious and non-infectious outcomes ^4^. School environmental factors have been shown to affect respiratory and cognitive health in school-aged children, with potential repercussions on future opportunities ^5^. *S. aureus*, and its methicillin-resistant form (MRSA), is an environmentally-hardy pathogen found widely in community environments, including schools ^6^. The environment plays an important role in *S. aureus* spread between humans and can serve as a reservoir for human exposure, especially in classrooms where children spend a significant portion of their day ^7^. Therefore, environmental exposure assessment is essential to protect this vulnerable population.

Despite its importance, there is no consensus on optimal protocol for indoor environmental sampling for *S. aureus* and MRSA, particularly in high-burden settings such as schools. Numerous sampling protocols have been employed for routine and post-outbreak environmental surveillance, ranging from swabs, dipslides, contact plates, and electrostatic wipes ^1,8–10^. Given *S. aureus*’ high prevalence in school environments, it is critical to utilize a sampling method that has a low detection limit (i.e. the method can detect bacterium at low concentrations distinguishable from zero) and high analytical sensitivity (i.e. the method can differentiate between two close measurements), resulting in high test accuracy.

This study sought to evaluate two sampling methods’ performance in a study within school environments. The first technique used a flocked swab for sample collection, followed by direct plating for colony-forming unit (CFU) enumeration, which is a commonly performed sampling procedure. We compared this approach to a technique that uses an autoclaved electrostatic cloth collection device, followed by plating the filtered sample for CFU enumeration ^11^. We tested both methods for their sensitivity to detect *S. aureus* presence, and for their ability to differentiate between heterogeneous samples based on CFU counts. Important in our assessment was their implementation in the field, which was measured by adherence to sampling protocol.

## Methods

### Sampling

Environmental samples were collected from nine mid-Atlantic inner-city schools from April to October 2016 as part of an ongoing study, which was approved by the Johns Hopkins University Institutional Review Board. School samples were collected from third and sixth grade classrooms and common spaces (cafeterias, gymnasiums, and libraries) throughout the year (spring, summer, fall). The location sampled was standardized for each room: two 31 cm^2^ (12 × 12 inches) target surface areas from adjacent but not overlapping areas on the primary door. Study personnel were trained to sample one area with a sterilized dry electrostatic cloth (Swiffer, Proctor & Gamble, Cincinnati OH), and the other area with a flocked swab (E-swab, Copan Diagnostics Murrieta CA), and alternated the collection technique used in the area directly near the door handle versus adjacent. The study team then measured and reported the actual area sampled, to test for any deviation in sampling protocol. Electrostatic cloths were placed into a sterile Stomacher blender bag (Fisher Scientific, Hampton NH) for transport, and the swabs were placed into Amies medium via the provided tube. Field staff used sterile collection methods, and a blank electrostatic cloth was obtained as a negative control for each visit to ensure sterile technique. Samples were stored in −4°C and processed within 72 hours of collection.

### Laboratory Processing

Samples were processed as reported in previous validation studies ^1,11^. Briefly, the Amies medium from the swabs was directly plated unto Baird-Parker Agar (Hardy Diagnostics, Santa Maria CA), incubated at 37°C for 48 hours, then coagulase positive colonies were counted. The electrostatic cloths were eluted with 50 ml phosphate-buffered saline, homogenized by hand, and filtered through 47-mm diameter, 0.45-µm pore-sized filter membranes (EMD Millipore, Billerica MA). The filter membrane was then placed unto a *Staphylococcus*-specific ChromAgar plate (ChromAgar™, Paris France), incubated at 37°C for 48 hours, then *S. aureus* phenotypic colonies were counted.

### Analysis

All statistical analysis was carried out on Stata statistical software (STATA 14, College Station, TX). Co-exposure variables included school, room type, and visit season. Outcomes were recorded as a continuous variable (exact CFUs) as well as dichotomous positive (1 or higher CFU count) or negative (0 CFU count). Continuous results were reported as total count CFU per sample, by extrapolating from the media volume used for culture to the total volume in the sample. We then standardized the total CFU count by the true surface area sampled, as reported by the study team. Test sensitivity was measured by the detection of *S. aureus* for each electrostatic cloths and swabs compared to positive for either technique. Linear and logistic regression models were run to control for confounding by school, room type, and season in the association of collection method with continuous CFU count and binary *S. aureus* detection.

## Results

Within the study time period from April to October 2016, 31 total surfaces were sampled with both swab and electrostatic cloth methods in 9 schools. **Table 1** provides a characteristics summary of the 31 samples obtained for analysis. As we were only able to sample for a select time period, we performed *post-hoc* power calculations, based on existing literature, to ensure adequate sample size. Based on previous literature of an expected prevalence of at least 50% positive and an estimated test sensitivity difference of 35% ^6,10^, we calculated that we would need a sample size of at least 27 sites per sampling technique (54 total samples) to achieve 80% or greater power to detect the expected difference in test sensitivity (Pearson’s chi-squared test) – less than our obtained 31 sites.

**Table 1:** Characteristics of school surface samples included in analysis

| Sample Characteristics | Count (Percent) |
| --- | --- |
| Unique Schools | 9 |
| Number of surface sample sites | 31 |
| **Sample Locations** |  |
| Classroom – 3<sup>rd</sup> grade | 12 (38.7%) |
| Classroom – 6<sup>th</sup> grade | 7 (22.6%) |
| Cafeteria | 7 (22.6%) |
| Gym | 2 (6.4%) |
| Library | 3 (9.7%) |
| **Visit Season** |  |
| Spring | 7 (22.6%) |
| Summer | 6 (19.4%) |
| Fall | 18 (58%) |

*S. aureus* was detected (CFU ≥ 1) from 31 (100%) electrostatic cloth samples and from 9 (29%) swab samples, as shown in **Table 2**. Using any detection on either test as the gold standard, the electrostatic cloth method had a 100% sensitivity (95% CI 98.93 – 100%), while the swabs had a 29.03% sensitivity (95% CI 14.22 – 48.04%). None of the blank control electrostatic cloths captured *S. aureus*, suggesting minimal contamination and adherence to sterile technique.

**Table 2:** Electrostatic Cloths Versus Swabs Culture Results

| Sensitivity Analysis, count (analytical sensitivity) |  |  |  |
| --- | --- | --- | --- |
|  | Electrostatic Cloths Positive | Electrostatic Cloth Negative | Total |
| Swab Positive | 9 | 0 | 9 (29.03%) |
| Swab Negative | 22 | 0 | 22 |
| Total | 31 (100%) | 0 | 31 |
| Continuous CFU Count of Electrostatic Cloth vs. Swab Samples, total and standardized |  |  |  |
|  | Electrostatic Cloths | Swabs | Difference (Cloth-Swab) |
| **Surface Area (cm<sup>2</sup>)** |  |  |  |
| Mean | 804.63 | 510.72 | 293.91 ** |
| 95% CI | 695.56 – 913.7 | 366.69 – 654.75 | 170.05 – 417.78 |
| Median | 929.03 | 232.26 | 83.87 ## |
| IQR | 780.64 – 929.03 | 232.26 – 929.03 | 0 – 696.78 |
| **Total CFU Count per Sample** |  |  |  |
| Mean | 36.13 | 7.09 | 29.03 * |
| 95% CI | 15.61 – 56.65 | 0 – 14.43 | 6.77 – 51.29 |
| Median | 23 | 0 | 16 ## |
| IQR | 7 – 43 | 0 – 5 | 4 – 36 |
| **Total CFU Count per 100 cm<sup>2</sup> surface area** |  |  |  |
| Mean | 5.23 | 4.5 | 0.73 |
| 95% CI | 2.49 – 7.97 | 0 – 10.24 | -5.93 – 7.38 |
| Median | 2.48 | 0 | 1.85 # |
| IQR | 1.07 – 6.02 | 0 – 0.39 | 0.32 – 4.31 |

Paired t-test for difference in means \*\* $p < 0.001$ , \* $p < 0.01$
Wilcoxon matched-pairs signed-ranks test for difference in medians ## $p < 0.001$ , # $p < 0.01$
CI = confidence interval, IQR = interquartile range

For CFU count as a continuous variable, the data were right skewed. Therefore, the median and interquartile ranges, in addition to the means and 95% confidence intervals, are presented in **Table 2**. A greater reported mean surface area was sampled with the electrostatic cloths compared to the swabs (804.63cm^2^ vs. 510.72cm^2^ respectively). The differences in surface area sampled within locations was 293.91cm^2^ mean (95% confidence interval 170.05 – 417.78) and 83.87cm^2^ median (0 – 696.78 interquartile range) – with a p-value <0.001 for paired t-test and Wilcoxon matched-paired test for significant difference in mean and median respectively. Electrostatic cloths had a higher mean and median total CFU count compared to the swabs (mean 36.16 CFU vs 7.09 CFU, median 23 CFU vs 0 CFU, respectively). After standardization by sampled area, electrostatic cloths showed a higher mean and median CFU count than swabs (mean 5.23 per 100cm^2^ and 4.5 per 100cm^2^, median 2.48 CFU per 100cm^2^ and 0, respectively). No confounding was found between school, room type, or season with continuous CFU count using a linear regression model, or binary *S. aureus* detection with a logistic regression model, regardless of collection method utilized.

## Discussion

Because environmental *S. aureus*, and MRSA specifically, in classroom spaces can have detrimental effect on childhood health, our study evaluated two protocols for sampling *S. aureus* in the indoor environment. All the electrostatic cloth samples detected *Staphylococcus aureus* from the sampled surfaces, resulting in higher analytic sensitivity compared to the swab samples (29%). Other studies evaluating MRSA in classrooms and public spaces have identified environmental prevalence ranging from 3.87% to 32.56% ^12,13^. However, these studies all utilized the swab collection technique. Our swab sample prevalence (29%, or 9/31) is comparable to these previous studies, while our electrostatic cloth samples showed a significantly greater prevalence (100%), indicating that this technique may be optimal, and potentially useful in other environments.

In addition to more frequent detection of *S. aureus* in the environment, our electrostatic cloth samples produced greater distribution among samples, which is important for identifying differences between samples. The electrostatic cloths had a higher median overall CFU count and area-standardized CFU count than the swab samples, as well as greater interquartile ranges. This greater range from the electrostatic cloth samples allows better characterization of the heterogeneity between different room samples. Due to a lower limit of detection, the swab sample quantities were clustered together, making it more difficult to compare bacterial burden between sample sites. This can impair the ability to evaluate for risk or protective factors.

Limitations of the results are the small sample size and the variability in sampled areas between the two collection techniques. The goal of the sampling technique was to capture adjacent, non-overlapping spaces of the same surface area, with the assumption that the two spaces would have equal *S. aureus* contamination. However, in reality the two zones could have had slightly different bacterial concentrations on the surfaces based on their unique micro-environments. We assume these variances result in a non-differential bias and designed the sampling protocol to minimize differential bias by alternating the sampling technique used in those two adjacent spaces, in order to systematically divide the variability in micro-environments between sampling protocol groups.

While our data showed electrostatic cloths to be more sensitive, this may in part be due to technique, but may in large part reflect field implementation. One significant finding in our study was the variability in surface area sampled, despite having a standardized protocol. This deviation from protocol reflects the importance of considering the practicality and real-world sampling conditions in the field. It should be noted that this was a highly trained research team who worked on multiple environmental exposure assessment studies. Despite reinforcement, the study team consistently under-sampled the target 31 cm^2^ area with the swab technique. By not collecting a sufficient surface area, there is a risk of not adequately depicting bacterial burden in certain locations. From anecdotal reports from the field, research staff felt that it was more difficult to capture the target surface area with the smaller swab as opposed to the larger electrostatic cloths, which better covers the designated sampling area due to the actual size of the collection surface. It was also reported to be faster to collect samples with the electrostatic cloths compared to the smaller swabs, saving valuable field time. There is a tradeoff in that the use of the electrostatic cloth technique requires staff training in sterile sample collection, but once learned, does not add significant time to sampling protocol. Further research would benefit from systematically capturing research staff opinions, in order to best design optimal study methodology from the perspective of barriers and facilitators to field staff protocol adherence.

This study demonstrated that the feasibility of a sampling technique in practice is as important as the laboratory efficiency. In this study, electrostatic cloths were shown to be more accurate, as well as practical in the field, for environmental *S. aureus* sampling. Based on the high sensitivity and differentiation among samples, this technique can be well adapted to the sampling of multiple environments and has been evaluated in indoor home sampling. Considering the importance of indoor environmental sampling in the context of human health outcomes, study methodology should be designed with significant consideration to both the efficacy and feasibility of sampling protocols for exposure assessment.

## Acknowledgements

The authors wish to thank Baltimore City Public Schools for their contributions. The authors would like to credit the following study team members for their collaboration – Larry Meade, Andres Lam, Fred Norton, Ehsan Majd, Christine Gummerson, Andrea Christ, and Ayanna Crear.

